# END-TO-END CLASSIFICATION OF CELL-CYCLE STAGES WITH CENTER-CELL FOCUS TRACKER USING RECURRENT NEURAL NETWORKS

**DOI:** 10.1101/2022.11.01.514198

**Authors:** Abin Jose, Rijo Roy, Dennis Eschweiler, Ina Laube, Reza Azad, Daniel Moreno-Andrés, Johannes Stegmaier

## Abstract

Cell division, or mitosis, guarantees the accurate inheritance of the genomic information kept in the cell nucleus. Malfunctions in this process cause a threat to the health and life of the organism, including cancer and other manifold diseases. It is therefore crucial to study in detail the cell-cycle in general and mitosis in particular. Consequently, a large number of manual and semi-automated time-lapse microscopy image analyses of mitosis have been carried out in recent years. In this paper, we propose a method for automatic detection of cell-cycle stages using a recurrent neural network (RNN). An end-to-end model with center-cell focus tracker loss, and classification loss is trained. The evaluation was conducted on two time-series datasets, with 6-stages and 3-stages of cell splitting labeled. The frame-to-frame accuracy was calculated and precision, recall, and F1-Score were measured for each cell-cycle stage. We also visualized the learned feature space. Image reconstruction from the center-cell focus module was performed which shows that the network was able to focus on the center-cell and classify it simultaneously. Our experiments validate the superior performance of the proposed network compared to a classifier baseline.

## 1. INTRODUCTION

Cell division is a relatively short but crucial and very dynamic period of the cell-cycle [1]. During mitosis, the cytology complexity reaches its superb zenith as eukaryotic cells rearrange structurally and functionally to accurately segregate each of the two replicated copies of the genetic material into two daughter cells. Errors in cell division are considered major sources of genomic instability, and have pathological consequences associated with diseases, including cancer formation, progression, and metastasis [2]. Therefore, the analysis of the cell-cycle, is paramount to accurately identify and characterize pathological phenotypes in clinically relevant situations. According with the traditional cytology of the cell nucleus [3, 4] cell-cycle is divided into interphase and mitosis, and the last phase into the sub-phases, prophase, prometaphase, metaphase, anaphase, and telophase. Quantification of the phase duration and topological characteristics of the cell nucleus and other structures is therefore crucial for determining the effect of drug treatments [5, 6, 7, 8]. The state-of-the-art approaches extracts hand-engineered features and then classify them using clustering algorithms. These methods do not use deep learning techniques. Zhong et al. [9], in 2012, proposed an unsupervised method for identifying the stages of mitosis. They introduced a clustering algorithm based on a temporally constrained combinatorial clustering (TC3) method. This approach uses the features extracted from the time-lapse microscopy images of human tissue culture cells using CellCognition [10] software. Flow cytometry is another advancement in the study of cellcycle [11], but it lacks the temporal component of cell-cycle progression, especially in the short time of mitotic sub-phases, potentially missing valuable information. According to [12, 13] DNA content could be used for identifying the phases of the cell. These methods rely on perfect segmentation of the cell nucleus for accurate identification. Feature extraction from fluorescence images followed by K-means clustering was performed by Ferro et al. [14] for classification. Conventional image processing methods extract the features and this feature selection in general affects the classification accuracy.

### 1.1. Related work

In recent years, deep learning methods [15, 16, 17], to name a few, were widely used for feature extraction and classification. For instance, Narotamo et al. [18] investigated three deep neural network-based learning approaches for classifying interphase cellcycle stages in microscopy images. Their main goal was to classify each DAPI-labelled nucleus from microscopic images. Another approach was proposed by Jin et al. [19] for cell-cycle image classification based on WGAN-GP and ResNet, and they tried to address the data imbalance problem with the cell-cycle detection datasets. Although, deep learning approaches have been used to detect mitosis in fixed tissue samples [20, 21, 22], it has not been used extensively in video sequences of cells undergoing mitosis. CellCognition [10] is a software tool that is applied for annotating time-resolved mitosis stages from sequences. The high similarity between images in different stages of mitosis will introduce high classification noise at the state transitions. The authors demonstrate that the incorporation of time information into the annotation can reduce this noise. They combined a supervised machine learning technique with a hidden markov model (HMM) [23] for classification.

### 1.2. This paper

We propose a method to identify different stages of mitosis using a deep learning technique by incorporating time information. We propose a novel RNN-based architecture to incorporate time-related propagation of features for classification. The main contributions of this paper are: 1) The proposed network is shallow and is trainable end-to-end. 2) The approach has a center-cell focus tracker module, to eliminate background information from other cells and to track the features of the cell undergoing mitosis. 3) A classifier to identify the cell stages. 4) Our approach incorporates the time information in the feature space which is inherent to the RNN. The main challenge is to train both the tracker and classifier together. We also compare the performance of proposed RNN-based approaches to the deep learning-based classification. The visualization of the learned feature space shows that the proposed model follows a time trajectory, unlike the classifier which forms individual clusters.

## 2 PROPOSED APPROACH

In 2016, Ondruska et al. [24] proposed an RNN-based approach for the end-to-end detection of objects from sensor data. Since in RNN the modules are connected in time, it can effectively transfer information across temporal data. The network can perform a semantic classification from the feature information learned in the hidden states. Ondruska et al. proposed that this network can be trained endto-end with a small amount of labelled data. A classification network is introduced once the deep tracking network is learned. To be able to track moving objects throughout the time sequence, the network must remember the location and other properties of each object. To achieve this tracking, gated recurrent units (GRUs) are used as the processing steps at each layer of the RNN. From the learned hidden layer, two convolutional networks are employed to predict the semantic segmentation as well as the mitosis class. Inspired by this model, we designed a novel architecture that consists of four main parts: 1) a backbone network for feature extraction, 2) a time encoding network to incorporate time information with the features, 3) a center-cell tracking network to focus the center-cell, and 4) a classification network for the mitosis stage prediction. The proposed architecture is illustrated in Fig. 1. This model is trained end-to-end by giving a cell video sequence as input. The tracking network and the classification network are trained simultaneously. Each layer of the proposed architecture is explained in detail below.

**Fig. 1:**
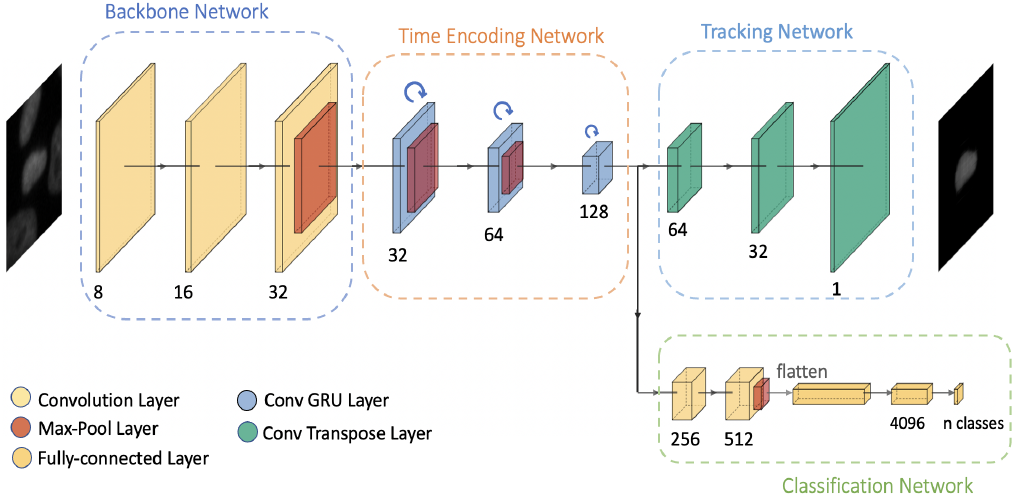
Illustration of the architecture of our proposed model applied to *n*^*th*^ frame of a video sequence with the four main parts.

### 1. Backbone network

This model uses a shallow backbone network and extracts high-dimensional features. It consists of three convolutional layers and a max-pooling layer. All three convolution layers use 3*×*3 kernels with the same padding and no striding. This network uses 8, 16, and 32 channels in each of the convolutional layers. Starting with the initial image 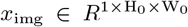 (grayscale image), the convolutional backbone generates a lower resolution activation map, *f ∈ R*^C*×*H*×*W^. The empirical values used for channels and image resolution are C = 32 and, 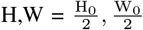 respectively.

### 2. Time encoding network

The time encoding network combines the extracted features from the previous frame to the current frame. Convolutional GRUs are the type of RNN employed in this work to combine the features between frames. The output of each GRU unit is given by a weighted combination of the current input, *n* and the previous output of frame *n −* 1. In this network, three convolutional GRUs with different channel sizes and dimensions are used to encode time information at different scales. 3 *×*3 kernels are used in these convolutional GRUs. A 2*×*2 max-pooling operation reduces the dimensionality of features by half after each GRU layer. 32, 64, and 128 respectively are the output channel sizes of each convolutional GRU layer. For the input size 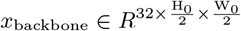, the typical output of our time encoding network has a channel size of 128 and image dimensions, 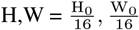 respectively.

### 3. Tracking network

An image frame may contain multiple cells. The main aim of our tracking network is to focus the features on the center-cell. The tracking network acts as a decoding network to reconstruct the center-cell from the output of the time encoding network.The time encoding network outputs features at a lower dimension. Three convolutional transpose layers are used to reconstruct the features to the original image dimensions. The typical values of the channel dimensions used in convolutional transpose layers are 64, 32, and 1 respectively. Thus, a resolution equal to the grayscale input image is reconstructed and the output dimensions are 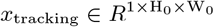, which has the same dimensions as *x*_img_.

### 4. Classification network

The classification network takes the output of the time encoding network as input features and then classifies these features into the number of required mitosis phases. The classification network consists of two convolutional layers, a max-pooling layer, and two fully-connected layers. The convolutional layers have channel sizes of 256 and 512. After the max-pooling reduces the dimensions, the features are flattened before passing through the fullyconnected layers. The size of the final layer of the fully-connected network is the number of mitosis states. This network predicts the class to which each input frame belongs. ^1 2^

## 3. TRAINING DETAILS

### 1. Training the tracking network

In the supervised training method, the ground truth data annotated by the experts is used to calculate the losses with the predicted outputs. The proposed model with the time encoding network has an additional tracking network to focus on the center-cell. Time encoding networks are used to help the flow of information across frames. Each image in the sequence data is dependent on previous images. A sequence of images belonging to the same cell is given as the input. The input images used for training the network are grayscale images with a resolution of 96*×*96 pixels. The tracking network reconstructs only the center-cell from the input image. This is achieved by calculating the loss (L_track_) between the predicted output and a masked image. The masked image is the input image masked with the segmentation of the center-cell. The last layer of the tracking network uses a sigmoid activation layer, as the images are normalized to the range [0, 1]. Then the loss between the predicted output and the masked image is calculated using a binary cross-entropy loss [25].

### 2. Training the classification network

This part of the network is trained with a one-hot vector of the ground-truth values. The training loss function used is binary cross-entropy loss (L_cls_). The total loss is calculated as a weighted sum of the tracking and classification losses. A regularization parameter *λ*_reg_ is used to combine the losses and the total loss is calculated as, L_tot_ = L_track_ + *λ*_reg_ *\** L_cls_.

## 4. EXPERIMENTAL SETUP

### The 3-class dataset

This dataset is provided by LiveCellMiner [26] and contains microscopic image sequences of the mitosis process. The LiveCellMiner tool allows the analysis of mitotic phases in 2D+t microscopy images. The dataset is named as LSM710 dataset [9] as the cells were acquired using an LSM710 confocal microscope (Zeiss) and ZEN software (Zeiss). These image sequences contain live human HeLa cells where the chromatin is labelled by the stable expression of histone 2 fused with the fluorescent protein mCherry. The acquired images have many cells in each time frame. The software [26] tracks the position of each cell in the image and then extracts the tracked single-cell image. LiveCellMiner then identifies and eliminates cells that do not undergo cell division. A region is cropped around each tracked cell from each time frame if the cell undergoes a cell division. Each of these cropped images has a resolution of 96*×*96 pixels, and the target cell is in the center of the cropped image. For each cell division sequence, 90 frames are available with this resolution. The dataset is divided into 3 phases such as interphase to early prophase as first state, late prophase to anaphase as next state, and telophase to interphase as the third state. The images were taken with a temporal resolution of three minutes. For training, 1458 video sequences were used.

### The 6-class dataset

The second dataset provided by Zhong et al. [9] also uses time-lapse microscopy images of human tissue culture cells (HeLa cells) expressing a fluorescent chromatin marker. Each frame in the dataset is labeled as interphase or one of the five mitosis stages. In the dataset, the labeling of interphase recovery is also included after cell division. The dataset consists of seven image sequences with a total of 326 cell division events uniformly sampled with an interval of 4.6 minutes. Since mitosis stages occur less frequently compared to the interphase stage, the distribution of cells in different classes is uneven. In this dataset, each image has a resolution of 96*×*96 pixels, and each cell image sequence has a length of 40 frames.

### Training the ResNet baseline model

To compare the performance of the proposed approach, we trained a ResNet18 [27] model as the classifier proposed in [19]. Here we consider each frame in the video as independent of other frames in the same sequence, without incorporating time information. A pre-trained model with the last layer replaced with the number of classes is used. Then the complete network is trained with images of mitosis sequence, with classification loss (L_cls_). The final layer of this network has the number of outputs equal to the number of mitosis phases to predict. A one-hot encoding vector of the ground truth is used to train this network. This network is trained with cross-entropy loss [25] to predict the probability of the image belonging to each class.

### Evaluation criteria and hyperparameters

We have calculated frame-to-frame accuracy from each sequence and averaged it over the total length of the test data. We have also evaluated the precision, recall, and F1-Score. The batch size equals the number of sequences. Due to the graphics unit’s memory limitations, deeper networks cannot employ batch sizes greater than 4 and hence we have selected 4 as the batch size. For the ResNet18 model, we have used a batch size of 64. The learning rate used is 0.01 for the proposed model and 0.001 for the ResNet18 model. The loss regularization value for the time encoded network is selected as 0.01. The train test ratio is chosen as 0.85. The learning rate is scheduled to reduce to 10 percent of the current value after 2500 iterations. Besides these hyperparameters, data augmentation is introduced in all our experiments to increase the robustness. Rotations by multiples of 90 degrees and flips in vertical and horizontal directions are the augmentations used. The training is conducted for 40 epochs for the 6-class dataset and 12 epochs for the 3-class dataset. Adagrad optimizer is used for training.

## 5. EXPERIMENTAL RESULTS

### Results on the 3-class dataset

We plotted the label matrix in which the y-axis represents different cell trajectories, and the x-axis represents the length of each trajectory. Fig. 2 (a) shows the label matrix of the ground truth annotation and the labels generated by the proposed network, on 50 test sequences. Green, magenta, and red represent the interphase class, mitosis class, and post-mitosis class respectively. On close examination, it is clear that the proposed model has results similar to user annotation. This is further substantiated by the classification accuracy of the proposed model which corresponds to 99.54%. We have better classification performance compared to the ResNet18 classifier which gave 99.30% accuracy. This means that introducing features from the previous time step using GRU layers helps in increasing the performance. The precision, recall, and F1-Score on the LSM710 dataset are shown in Table 1. It is evident from Table 1 that using the time-encoded network improves the performance compared to the classification model. Even though the precision and recall values of the classification model got comparable results, our proposed model has higher F1-Score values for three different cell division phases.

**Table 1:** Precision, Recall and F1-Score for the 3-class dataset.

| Method | Precision |  |  |
| --- | --- | --- | --- |
|  | Phase1 | Phase2 | Phase3 |
| Ours | 98.649 | **97.988** | **99.976** |
| ResNet [19] | **99.081** | 97.658 | 99.686 |
|  | Recall |  |  |
|  | Phase1 | Phase2 | Phase3 |
| Ours | **99.806** | **97.872** | 99.617 |
| ResNet [19] | 99.179 | 96.927 | **99.801** |
|  | F1-Score |  |  |
|  | Phase1 | Phase2 | Phase3 |
| Ours | **99.259** | **97.930** | **99.797** |
| ResNet [19] | 99.130 | 97.291 | 99.744 |

**Fig. 2:**
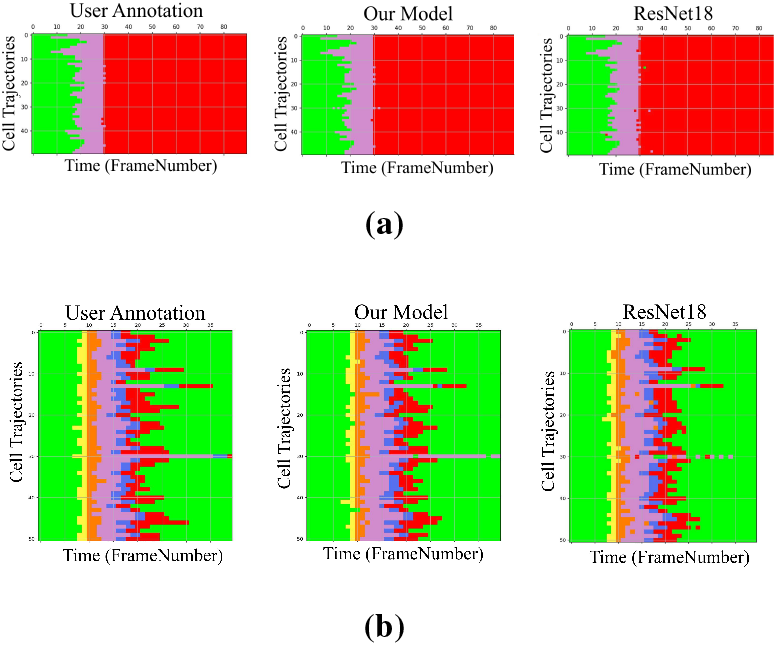
Label matrices for the user annotation and the predicted labels by the network for the proposed model, and the ResNet18 classifier on test sequences for (a) the 3-class dataset, (b) the 6-class dataset. The color code corresponds to the cell-cycle phases.

Fig. 4 shows the reconstructed output image from the tracking network at three different time points of the video sequence. The tracking network is able to focus on the center region of the frame, which corresponds to the cell under consideration and has eliminated the information from the surrounding cells. The embedding space gives characteristics related to some properties of the input data. Fig. 6 (a,b) illustrates the first three principal components of the embedding space of our proposed model. In theory, during the postmitotic stage, the daughter cells will grow back to the interphase stage, called interphase recovery. Since the original ResNet18 operates on individual images instead of sequences, the overlap between interphase and post-mitosis features happens because of this interphase recovery. This does not occur in network with GRU layers because the time information is involved which in turn is the main advantage of our approach. This is also evident in the embedding space plot as the features are connected and move from blue-red-pink classes corresponding to the three mitosis phases. Whereas the embedding plot corresponding to the classifier forms three clusters and due to interphase recovery, the blue and pink classes are having overlap resulting in miss-classification.

### Results on the 6-class dataset

This section compares the performance of the proposed network with the morphology dataset by Zhong et al. Fig. 2 (b), depicts the label matrix of the ground truth annotation and the labels generated by the user annotation, and the networks on test sequences. The y-axis in this plot represents the 51 test sequences, and the x-axis is the length of each cell sequence. Each cell sequence in this dataset has 40 frames. The predicted label matrix of the proposed model compared to the user annotation labels looks comparable. This indicates that the network was able to predict closer to ground truth annotation. A more precise evaluation can be done by analyzing the frame-to-frame accuracy of each model. Our proposed model has a higher frame-to-frame accuracy of 93.137%, while the ResNet18 classifier has a lower accuracy of 92.254%.

Table 2 shows the precision, recall, and F1-Score of the proposed model and classifier on each of the 6 phases. The results from our proposed network architecture are compared with the ResNet classifier. Our model has similar or slightly higher performance compared to ResNet classifier, except for the prophase. The lower number on prophase is due to the confusion between the last frames of interphase and prophase frames. It is clear from the confusion matrix shown in Fig. 3. Most of the prophase images are misclassified as interphase images. This is also evident in the reconstruction results shown in Fig. 5 (g,h). The main reason for this confusion is that interphase images are just one or two frames longer and is an underrepresented class and are mostly closer in appearance to the last frames of interphase class. The embedding space visualization is illustrated in Fig. 6 (c,d) and has similar behavior to the 3-class dataset. Since the cells begin and end at interphase (blue), the time encoded network has an embedding space with classes moving from one state to the next which begins and ends in interphase. However, the classifier forms 6 different clusters in the embedding space. Fig. 5 shows the reconstruction results of the tracker module for images in 6 different phases. This clearly shows that the center-cell is reconstructed by the network eliminating clutter from the surrounding cells.

**Table 2:** Precision, Recall and F1-Score for the 6-class dataset.

| Method | Precision |  |  |  |  |  |
| --- | --- | --- | --- | --- | --- | --- |
|  | Inter | Pro | Prometa | Meta | Ana | Telo |
| Ours | **95.45** | 82.81 | **92.63** | 89.68 | **90.20** | **88.00** |
| ResNet [19] | 95.18 | **88.57** | 87.50 | **92.69** | 85.22 | 80.88 |
|  | Recall |  |  |  |  |  |
|  | Inter | Pro | Prometa | Meta | Ana | Telo |
| Ours | **97.94** | 69.74 | 83.02 | **96.92** | 81.41 | **80.00** |
| ResNet [19] | 96.83 | **81.58** | **85.85** | 92.69 | **86.73** | 75.00 |
|  | F1-Score |  |  |  |  |  |
|  | Inter | Pro | Prometa | Meta | Ana | Telo |
| Ours | **96.68** | 75.71 | **87.56** | **93.16** | 85.58 | **83.81** |
| ResNet [19] | 96.00 | **84.93** | 86.66 | 92.69 | **85.96** | 77.83 |

**Fig. 3:**
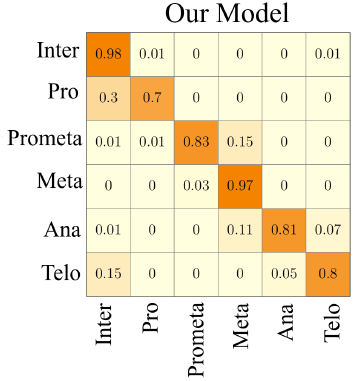
Confusion matrix for the 6-class dataset for the proposed approach.

**Fig. 4:**
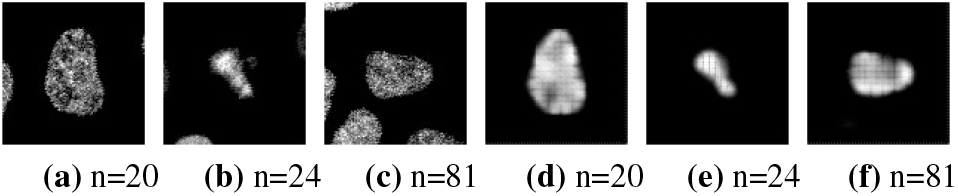
Input images (a-c) and output reconstruction (d-f) from the tracker at three different stages for the 3-class dataset. Frame numbers are denoted by *n*. Frames are captured three minutes apart.

**Fig. 5:**
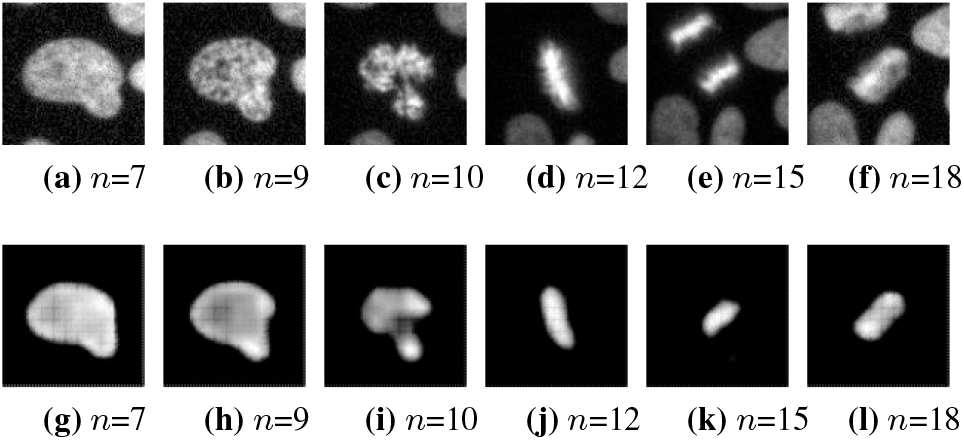
Input images (first row) and output reconstruction (second row) from the tracker at six different stages for the 6-class dataset. Frame numbers are denoted by *n*. Frames are captured 4.6 minutes apart.

**Fig. 6:**
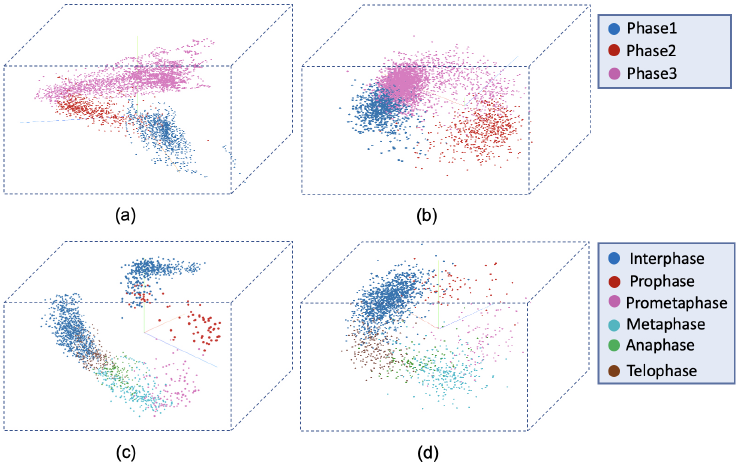
First 3 principal components of feature space for (a) proposed model on the 3-class dataset, (b) ResNet on the 3-class dataset, (c) proposed model on the 6-class dataset, (d) and ResNet on the 6-class dataset.

## 6. CONCLUSIONS

We proposed a novel deep learning-based approach for focusing on the center-cell in a video sequence and identifying the cell division stages in the sequence. We were able to achieve better classification performance using a shallower network with time information. The encoding of time information is achieved by RNNs and the network is trained end-to-end with a center-cell tracker and classifier loss. The feature space visualizations clearly show the advantage of using RNNs as the features are connected from one state to the next unlike the feature space of a normal classifier, which forms intermingled clusters. The evaluation of accuracy, precision, recall, and F1-Score gave higher numbers for the proposed model. In future, we plan to investigate the effect of deeper networks with time information to fine tune the miss-classifications in the 6-class dataset. Extension of network architecture to 3D datasets is another possible research direction.

## Footnotes

1 Code - https://github.com/Rijo756/cell-cycle-stages-identification.

2 Supplementary - https://www.lfb.rwth-aachen.de/wp-content/uploads/icassp.pdf.

## REFERENCES

[1] Ana Rita Araujo, Lendert Gelens, Rahuman SM Sheriff, and Silvia DM Santos, “Positive feedback keeps duration of mitosis temporally insulated from upstream cell-cycle events,” Molecular Cell, vol. 64, no. 2, pp. 362–375, 2016.

[2] Andréa E Tijhuis, Sarah C Johnson, and Sarah E McClelland, “The emerging links between chromosomal instability (cin), metastasis, inflammation and tumour immunity,” Molecular Cytogenetics, vol. 12, no. 1, pp. 1–21, 2019.

[3] Robert A Lambert, “Comparative studies upon cancer cells and normal cells: Ii. the character of growth in vitro with special reference to cell division.,” The Journal of Experimental Medicine, vol. 17, no. 5, pp. 499, 1913.

[4] Warren H Lewis and Margaret R Lewis, “The duration of the various phases of mitosis in the mesen chyme cells of tissue cultures,” The Anatomical Record, vol. 13, no. 6, pp. 359–367, 1917.

[5] Mojca Mattiazzi Usaj, Erin B Styles, Adrian J Verster, Helena Friesen, Charles Boone, and Brenda J Andrews, “High-content screening for quantitative cell biology,” Trends in Cell Biology, vol. 26, no. 8, pp. 598–611, 2016.

[6] Shuaizhang Li and Menghang Xia, “Review of high-content screening applications in toxicology,” Archives of Toxicology, vol. 93, no. 12, pp. 3387–3396, 2019.

[7] Srinivas Niranj Chandrasekaran, Hugo Ceulemans, Justin D Boyd, and Anne E Carpenter, “Image-based profiling for drug discovery: due for a machine-learning upgrade?,” Nature Reviews Drug Discovery, vol. 20, no. 2, pp. 145–159, 2021.

[8] Nikolaus Rajewsky, Geneviève Almouzni, Stanislaw A Gorski, Stein Aerts, Ido Amit, Michela G Bertero, Christoph Bock, Annelien L Bredenoord, Giacomo Cavalli, Susanna Chiocca, et al., “Lifetime and improving european healthcare through cell-based interceptive medicine,” Nature, vol. 587, no. 7834, pp. 377–386, 2020.

[9] Qing Zhong, Alberto Giovanni Busetto, Juan P Fededa, Joachim M Buhmann, and Daniel W Gerlich, “Unsupervised modeling of cell morphology dynamics for time-lapse microscopy,” Nature Methods, vol. 9, no. 7, pp. 711–713, 2012.

[10] Michael Held, Michael HA Schmitz, Bernd Fischer, Thomas Walter, Beate Neumann, Michael H Olma, Matthias Peter, Jan Ellenberg, and Daniel W Gerlich, “Cellcognition: time-resolved phenotype annotation in high-throughput live cell imaging,” Nature Methods, vol. 7, no. 9, pp. 747–754, 2010.

[11] Joe W Gray and Zbigniew Darzynkiewicz, Techniques in cell cycle analysis, Springer Science & Business Media, 1987.

[12] Vassilis Roukos, Gianluca Pegoraro, Ty C Voss, and Tom Misteli, “Cell cycle staging of individual cells by fluorescence microscopy,” Nature Protocols, vol. 10, no. 2, pp. 334–348, 2015.

[13] Damian J Matuszewski, Ida-Maria Sintorn, Jordi Carreras Puigvert, and Carolina Wählby, “Comparison of flow cytometry and image-based screening for cell cycle analysis,” in International Conference on Image Analysis and Recognition. Springer, 2016, pp. 623–630.

[14] Anabela Ferro, Tânia Mestre, Patrícia Carneiro, Ivan Sahumbaiev, Raquel Seruca, and João M Sanches, “Blue intensity matters for cell cycle profiling in fluorescence dapi-stained images,” Laboratory Investigation, vol. 97, no. 5, pp. 615–625, 2017.

[15] SanaUllah Khan, Naveed Islam, Zahoor Jan, Ikram Ud Din, and Joel JP C Rodrigues, “A novel deep learning based framework for the detection and classification of breast cancer using transfer learning,” Pattern Recognition Letters, vol. 125, pp. 1–6, 2019.

[16] Flávio HD Araújo, Romuere RV Silva, Daniela M Ushizima, Mariana T Rezende, Cláudia M Carneiro, Andrea G Campos Bianchi, and Fátima NS Medeiros, “Deep learning for cell image segmentation and ranking,” Computerized Medical Imaging and Graphics, vol. 72, pp. 13–21, 2019.

[17] Khalid Hamed S Allehaibi, Lukito Edi Nugroho, Lutfan Lazuardi, Anton Satria Prabuwono, Teddy Mantoro, et al., “Segmentation and classification of cervical cells using deep learning,” IEEE Access, vol. 7, pp. 116925–116941, 2019.

[18] Hemaxi Narotamo, Maria Sofia Fernandes, Ana Margarida Moreira, Soraia Melo, Raquel Seruca, Margarida Silveira, and João Miguel Sanches, “A machine learning approach for single cell interphase cell cycle staging,” Scientific Reports, vol. 11, no. 1, pp. 1–13, 2021.

[19] Xin Jin, Yuanwen Zou, and Zhongbing Huang, “An imbalanced image classification method for the cell cycle phase,” Information, vol. 12, no. 6, pp. 249, 2021.

[20] Anabia Sohail, Asifullah Khan, Noorul Wahab, Aneela Zameer, and Saranjam Khan, “A multi-phase deep cnn based mitosis detection framework for breast cancer histopathological images,” Scientific Reports, vol. 11, no. 1, pp. 1–18, 2021.

[21] Monjoy Saha, Chandan Chakraborty, and Daniel Racoceanu, “Efficient deep learning model for mitosis detection using breast histopathology images,” Computerized Medical Imaging and Graphics, vol. 64, pp. 29–40, 2018.

[22] Abdulkadir Albayrak and Gokhan Bilgin, “Mitosis detection using convolutional neural network based features,” in 2016 IEEE 17th International Symposium on Computational Intelligence and Informatics (CINTI). IEEE, 2016, pp. 000335– 000340.

[23] Greg M Gallardo, Fuxing Yang, Fiorenza Ianzini, Michael Mackey, and Milan Sonka, “Mitotic cell recognition with hidden markov models,” in Medical Imaging 2004: Visualization, Image-Guided Procedures, and Display. SPIE, 2004, vol. 5367, pp. 661–668.

[24] Peter Ondrúška and Ingmar Posner, “Deep tracking: Seeing beyond seeing using recurrent neural networks,” in Proceedings of the Thirtieth AAAI Conference on Artificial Intelligence, 2016, pp. 3361–3367.

[25] Christopher M Bishop and Nasser M Nasrabadi, Pattern Recognition and Machine Learning, vol. 4, Springer, 2006.

[26] Daniel Moreno-Andrés, Anuk Bhattacharyya, Anja Scheufen, and Johannes Stegmaier, “Livecellminer: A new tool to analyze mitotic progression,” PloS One, vol. 17, no. 7, pp. e0270923, 2022.

[27] Kaiming He, Xiangyu Zhang, Shaoqing Ren, and Jian Sun, “Deep residual learning for image recognition,” in Proceedings of the IEEE Conference on Computer Vision and Pattern Recognition, 2016, pp. 770–778.

